# Lactate promotes IL-8 secretion in human alveolar macrophages through GPR132 and lipid metabolic reprogramming

**DOI:** 10.64898/2026.01.27.701994

**Authors:** Myah Ali, Michelangelo Certo, Joseph Johnson, Atrayee Gope, Lauren Davis, Jodie Harte, Haleemah Hussain, Charlie Owen-Melody, Sally Clayton, Lucy Adam, Shannon L O’Brien, Davide Calebiro, Robert Lane, Babu Naidu, David Thickett, Aaron Scott, Claudio Mauro, Alba Llibre

## Abstract

Alveolar macrophages are the primary lung immune cell. They play a crucial role in both maintenance of tissue homeostasis and the initiation of inflammation, secreting multiple immune mediators including the chemokine interleukin-8 (IL-8). Inflammatory settings are often characterised by tissue hypoxia, increased glycolytic rates and lactate secretion, yet how lactate influences alveolar macrophage function remains unclear. Here we investigated how lactate, once considered a waste-product, shapes alveolar macrophage function. To do so, primary human alveolar macrophages (hAMs) and a combination of flow cytometry, ELISA/Luminex, western blot, light microscopy, bioluminescence resonance energy transfer (BRET) and analysis of publicly available datasets were used to understand the role of lactate in lung pathological environments. We demonstrate that hAMs sense extracellular lactate via the expression of different lactate transporters (e.g. MCT1, MCT4) and receptors (i.e. GPR132). Lactate treatment of hAMs increased IL-8 secretion in a MCT1-dependent manner. Lipid metabolism and lipid droplet formation, as well as direct lactate-driven GPR132 signalling were required for lactate-dependent IL-8 release. Our findings uncover a previously unrecognised dual mechanism by which lactate orchestrates immune regulation in hAMs. Specifically, lactate-driven IL-8 production requires two distinct lactate-driven processes: uptake via MCT1, which reprograms lipid metabolism, and signaling through the lactate receptor GPR132. This functional integration of lactate transport and receptor-mediated signaling provides new mechanistic insight into lactate’s role in hAM biology and highlights potential targets for therapeutic intervention.

## INTRODUCTION

Lung diseases are a global health concern, being major causes of disability and death worldwide^1^. The lung is a unique organ, characterised by being lipid-rich and glucose-poor ^2^. Through breathing and gas exchange, the lungs are constantly exposed to the outside world, thus confronted with a range of stimuli, mostly harmless. Alveolar macrophages (AMs) are the primary immune cell in the human lung. Because of the uniqueness of the physiological niche they occupy, AMs are primarily immune-suppressive at baseline, ensuring immune tolerance to innocuous inhaled particles^3^. Here, AMs guard lung homeostasis by phagocytosing debris (e.g. inhaled particles, apoptotic cells, surfactant) and releasing anti-inflammatory cytokines such as TGFβ and IL-10. When facing a challenge, AMs play a crucial role in initiating the inflammatory response. From pathogen detection and phagocytosis to secretion of pro-inflammatory mediators (e.g. IL-8, IL-6, IL-1β), they are essential in raising the alarm and orchestrating the initiation of an appropriate immune response. Specifically, IL-8 secreted by AMs act as potent neutrophil chemoattractant^4,5^. Neutrophils are a double-edge sword, being essential in the innate immune responses against pathogens, but also drivers of immune-pathology and tissue destruction if not fine-tuned^6^.

When homeostasis is broken, the cellular equilibrium is disrupted, causing a plethora of physiological changes, including metabolic adaptations of the cell types involved. This is in response to fluctuations in concentrations and/or availability of specific molecules, including glucose, fatty acids, amino acids and oxygen^7^. In turn, the cellular metabolic reprogramming - the reliance of cells on specific metabolic pathways - shapes both the intracellular and extracellular microenvironment^8^. The metabolic reprogramming observed in several lung diseases often results in lipid accumulation within AMs^9–11^. These fat-rich, foamy macrophages are a feature of several respiratory conditions, including chronic obstructive pulmonary disease (COPD) and tuberculosis (TB)^12^. Specifically, *Mycobacterium tuberculosis* (*Mtb*) reprograms cell metabolism promoting this lipid-laden macrophage phenotype. There is evidence suggesting this lipid accumulation might be both beneficial for the pathogen (e.g. access to nutrients) and the host (e.g. production of immune-modulatory lipid mediators)^13–15^.

In the lung, enhanced glycolysis and lactate production have been described in a range of pathologies. These include lung cancer, acute respiratory distress syndrome (ARDS), asthma, pulmonary hypertension (PH), idiopathic pulmonary fibrosis (IPF), COPD, as well as bacterial and viral infections (reviewed in Jia *et al.* 2025^16^). Often linked to hypoxia, these increased glycolytic rates appear to profoundly shape the local immune response^8,17,18^. Specifically, lactate is now recognised as a multifaceted immune-regulatory molecule^19^. Not only is lactate the primary gluconeogenic precursor and a key source for mitochondrial respiration, but it is also a multifunctional signalling molecule^19^. It can be imported from the extracellular milieu into the cytoplasm, and vice versa, depending on the concentration gradient. Specific lactate monocarboxylate transporters (MCTs) and sodium-coupled monocarboxylate transporters have been identified in different cell types^20^. Lactate can also directly signal through specific G protein-coupled receptors (GPCRs), triggering intracellular signalling cascades and impacting cell behaviour. For instance, GPR81 is mainly expressed in adipose tissue and lactate signalling through this receptor inhibits lipolysis^21,22^. In contrast, GPR132 is more broadly expressed and, when triggered by lactate, promotes an anti-inflammatory macrophage phenotype in the tumour microenvironment^23,24^. Inside the cell, lactate can be converted into pyruvate and fuel the TCA cycle. Its carbons can also contribute to the Acetyl-CoA pool^25–27^. Acetyl-CoA can be both used for energy production and as a building block for the synthesis of complex biomolecules, including fatty acids^28^. Thus, from carbon source to signalling molecule, lactate contributes to shaping cell metabolism and function. This study aims to provide insights into how human alveolar macrophages (hAMs) sense and respond to lactate, and what are the underpinning mechanisms of lactate-driven functional changes in this unique cell type.

## RESULTS

### Lactate promotes IL-8 secretion in alveolar macrophages in an MCT1-dependent manner

Glycolytic engagement is a common feature of inflammatory responses, with high glycolytic rates and increased lactate production described in various lung pathologies^16^. Given the key role of hAMs in maintaining lung homeostasis, we wanted to explore whether hAMs are equipped to sense and respond to environmental fluctuations of in lactate. We first confirmed that hAMs indeed express the lactate transporters SLC16A1 (MCT1) and SLC16A3 (MCT4) (Figure 1A and Supplementary Figure 1), indicating their capacity for lactate import and intracellular activity. To explore the impact of elevated lactate concentrations relevant to inflamed lung environments within the hAM population^29–35^, hAMs were exposed to 30mM lactate for 48 hours. Cytokine analysis of supernatants revealed increased secretion of pro-inflammatory mediators (e.g. MIP-1a, IL-8, IL-6) compared to untreated controls (Figure 1B). In inflammatory settings, circulating monocytes are recruited to the lung where they differentiate into macrophages^36^. To mimic these *in vivo* settings, we also differentiated human monocytes in the presence of 30mM lactate and observed increased release of pro-inflammatory cytokines (e.g. MIP-1b, MIP-1a, IL-8, IL-6) and decreased secretion of anti-inflammatory mediators (IL-1ra) (Figure 1B). Because IL-8 is a major orchestrator of inflammation and it often contributes significantly to disease pathology^37,38^, we further investigated how lactate triggers its secretion. Lactate promoted IL-8 secretion in a dose-dependent manner (Figure 1C, 1D), and this effect was dependent on lactate import through the MCT1 transporter (Figure 1E). Collectively, these data establish lactate as an immune regulator of human lung alveolar macrophages, promoting a pro-inflammatory phenotype.

**Figure 1.**
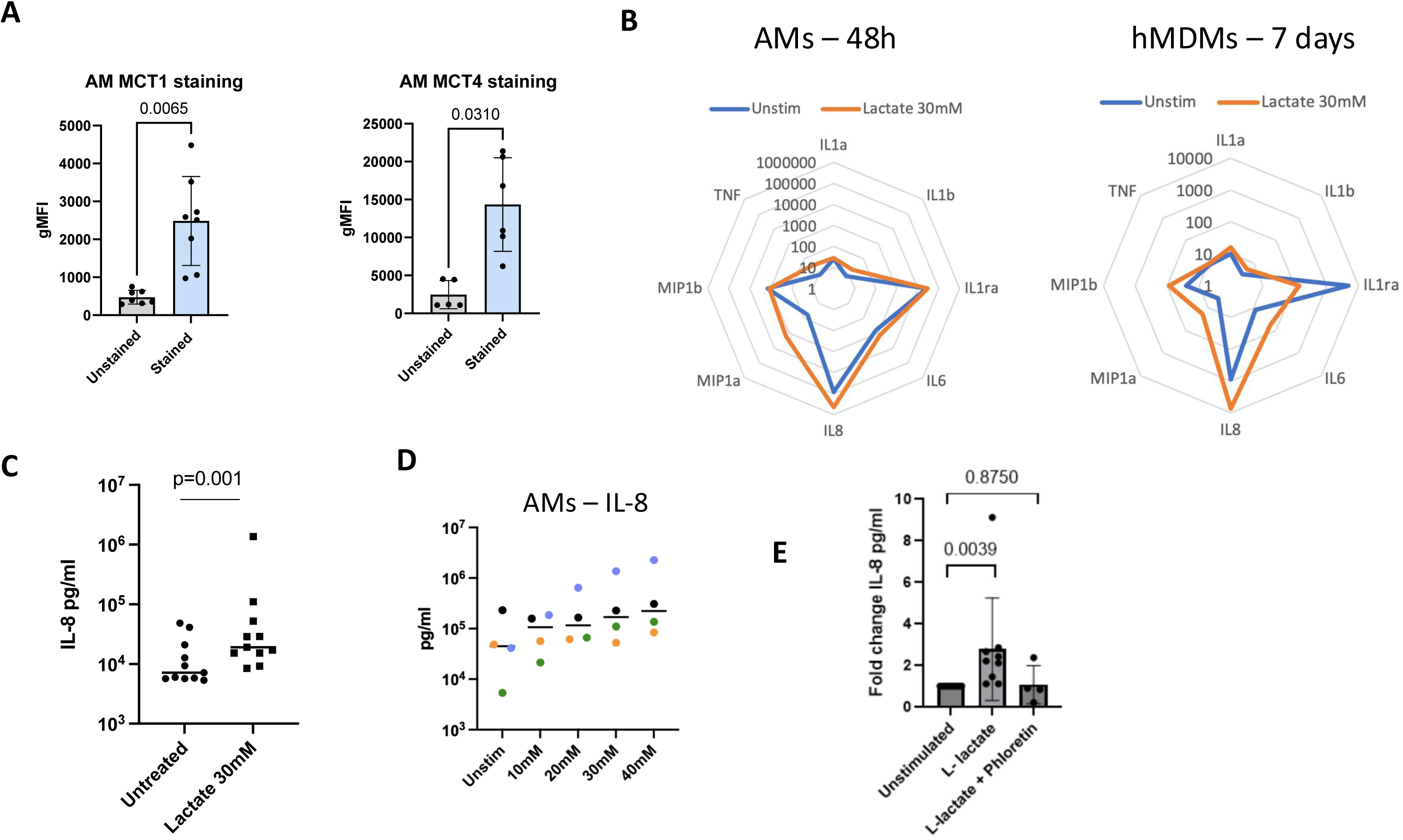
Lactate triggers IL-8 secretion in alveolar macrophages in a MCT1-dependent manner. **(A)** Expression levels of MCT1 (SLC16A1) and MCT4 (SLC16A3) in human alveolar macrophages (hAMs), measured by flow cytometry (n=7). **(B)** Cytokine concentrations, measured by Luminex (pg/ml, log scale), of supernatants from hMDMs differentiated in the presence of Na-lactate (30mM) (7 days, n=4) and hAMs treated with Na-lactate (30mM) for 48h (n=4). Supernatant concentration of IL-8, measured by ELISA, of hAMs treated with **(C)** Na-lactate (30mM) for 48h (n=10), **(D)** increasing concentrations of Na-lactate (n=4) or **(E)** Na-lactate (30mM) for 48h in the presence or absence of the MCT1 inhibitor phloretin (100μg/ml) (n=4).

### Lactate rewires lipid metabolism in alveolar macrophages, which is required for IL-8 secretion

Lactate-derived carbons fuel fatty acid (FA) synthesis^27,39–41^, with FAs serving as the building blocks for neutral lipid synthesis and the formation of lipid droplets (LDs). To assess the impact of lactate exposure on lipid content in hAMs, cells were treated with lactate for 48 hours and neutral lipid accumulation was assessed by Oil Red O (ORO) staining. Lactate treatment resulted in increased lipid content within hAMs, comparable to the positive control oleic acid treatment^42,43^ (Figure 2A). Mechanistically, lactate promotes FA and LD formation by providing carbons for acetyl-CoA synthesis^25–27^. Acetyl-CoA carboxylase (ACC) catalyses the rate-limiting step in fatty acid synthesis, converting acetyl-CoA to malonyl-CoA, and its activity is inhibited by phosphorylation^44,45^. We observed that lactate treatment substantially decreased ACC phosphorylation (pACC) without impacting total ACC levels (Figure 2B), potentially enhancing ACC enzymatic activity. This provides novel mechanistic insight into how lactate triggers lipid synthesis and accumulation within hAMs, in addition to its already defined role as a metabolic substrate (Figure 2C). To further investigate the relationship between lactate-driven IL-8 secretion (Figure 1C) and lipid metabolism, hAMs were treated with lactate in the presence or absence of inhibitors targeting specific lipid metabolic pathways. LDs are composed of triacylglycerols and steryl esters^46^. Using A922500 to inhibit diacylglycerol acyltransferase 1 (DGAT-1), the enzyme responsible for the last step in triacylglycerol synthesis, resulted in decreased lactate-driven IL-8 secretion (Figure 2D). Similarly, direct inhibition of ACC using TOFA and inhibition of fatty acid synthase (FAS) using C75 also diminished lactate-dependent IL-8 secretion in hAMs (Figure 2E and 2F). Of note, the inhibitors also decreased IL-8 secretion in the absence of lactate, highlighting lipid synthesis as a fundamental regulator of hAM pro-inflammatory activity. Taken together, these data demonstrate that increased lipid metabolism plays a key role in lactate-driven IL-8 secretion in hAMs.

**Figure 2.**
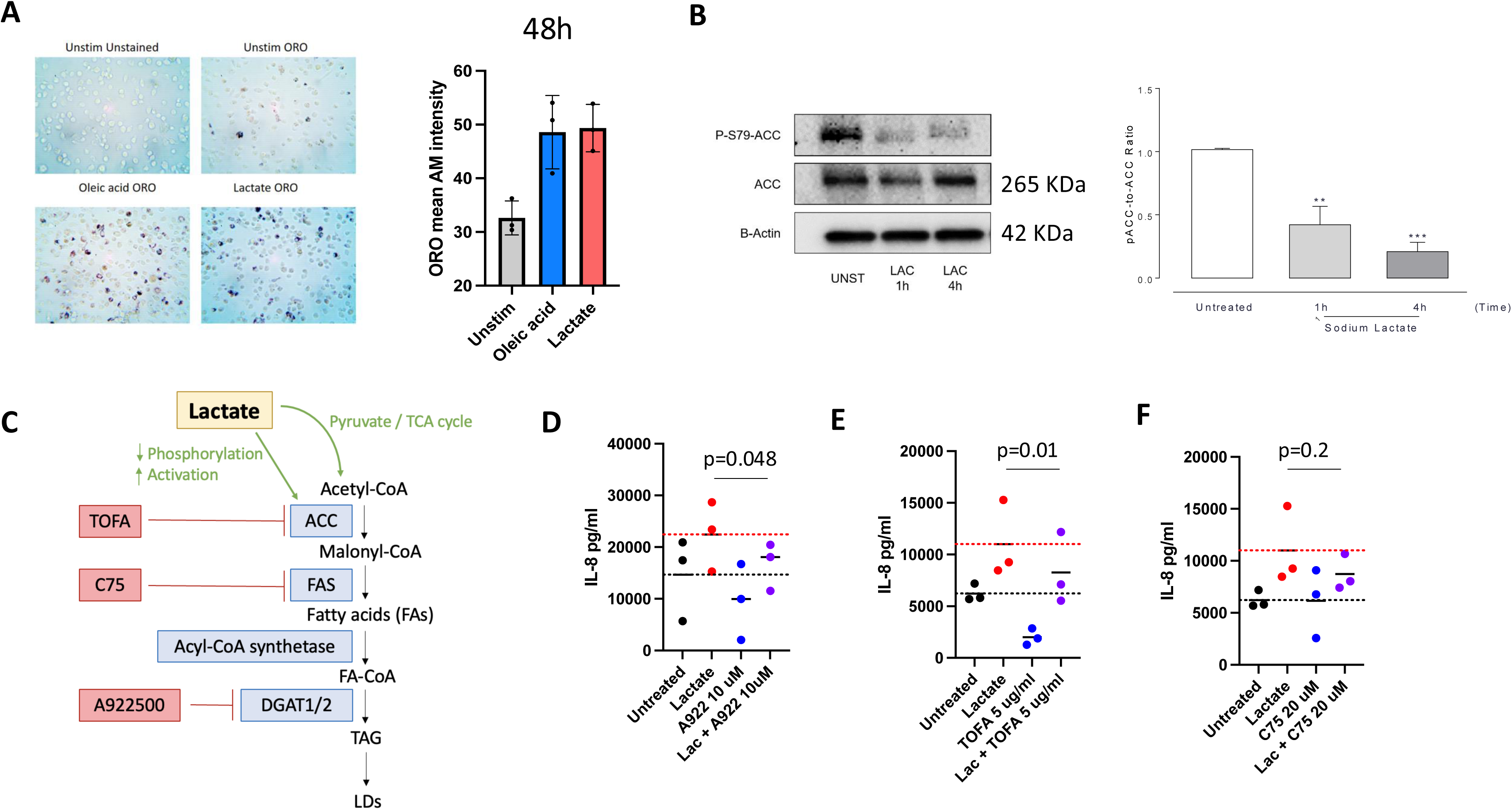
Lactate induces LD formation in alveolar macrophages, which is required for IL-8 secretion. Experiments performed in hAMs treated with Na-Lactate (30mM) for 48h, unless otherwise specified. **(A)** Oil Red O (ORO) lipid staining of AMs treated with oleic acid (Positive control; 30 μM) or Na-lactate. Representative picture (n=3). **(B)** Western blot and ratio of phosphorylated (deactivated) and unphosphorylated (activated) Acetyl-CoA carboxylase (ACC) of human AMs treated with lactate (1h and 4h, n=3). **(C)** Schematic of how lactate contributes to lipid accumulation and lipid droplet formation. ACC: Acetyl-CoA Carboxylase. FAS: Fatty Acid Synthase. DGAT: Diacylglycerol Acyltransferase. TCA: Tricarboxylic acid. FA: Fatty acids. TAG: Triacylglycerides. LDs: Lipid droplets. TOFA: 5-tetradecyloxy-2-furoic acid. C75: trans-4-carboxy-5-octyl-3-methylenebutyrolactone. A922500: DGAT-1 inhibitor. hAMs treated with Na-lactate +/− the indicated concentrations of **(D)** a LD inhibitor (A922500), **(E)** an Acetyl-CoA Carboxylase inhibitor (TOFA) to inhibit fatty acid synthesis, or **(F)** a Fatty Acid Synthase inhibitor (C75) and IL-8 measured by ELISA (n=3).

### *CXCL8* expression correlates with *PTGS2* and *GPR132* in hAMs infected with *Mtb*

Having demonstrated the importance of lipid metabolism for lactate-induced IL-8 secretion, the mechanisms of lactate-driven action were further investigated. LDs are well-known reservoirs of cyclooxygenase (COX) enzymes, also known as prostaglandin-endoperoxide synthase (PTGS)^47^. COX-1 and COX-2 synthesise prostaglandins (PGE), lipid mediators with important regulatory functions^48,49^. Specifically, PGE2 has been shown to promote IL-8 expression in several contexts and cell types^50–53^. Thus, it is plausible that lactate drives IL-8 secretion via elevation of PGE2. An alternative hypothesis is that lactate exerts its effects via direct signalling through specific GPCRs (i.e. GPR132 and GPR81)^54^ located at the cell surface and/or intracellularly^55,56^, particularly since MCT1 is key for lactate-driven IL-8 secretion (as illustrated in Figure 1E). These hypotheses were initially explored by investigating publicly available datasets. Specifically, RNAseq datasets from hAMs infected with *Mtb* were analysed^57–61^. *Mtb* infection is a suitable context for the testing of our hypotheses since 1) metabolic reprogramming of the lung and increased glycolysis upon *Mtb* infection is well-documented^62–64^, 2) lipid-rich, foamy macrophages are a hallmark of TB granuloma lesions^12^ and 3) IL-8 is produced by lung macrophages in the context of *Mtb* infection^65,66^. The data indicate that hAMs upregulated *CXCL8* (the gene encoding IL-8), *PTGS2* (which encodes COX-2) and *GPR132* expression, but not *HCAR1* (which encodes GPR81) after *Mtb* infection (Figure 3A). Moreover, *CXCL8* expression strongly correlated with *PTGS2* and *GPR132,* but not *HCAR1,* expression in both non-infected and *Mtb*-infected hAMs (Figure 3B, 3C and 3D). To test if this correlation was specific to *Mtb* infection, the Human Lung Cell Atlas (HLCA), an integrated single-cell reference atlas of the human lung^67^, was also interrogated. A similar trend was observed in both healthy and specific disease settings (pulmonary fibrosis, COPD) (Supplementary Figure 2). These data indicate that both COX enzymes and GPR132 may be involved as mediators of lactate-driven IL-8 secretion in hAMs.

**Figure 3.**
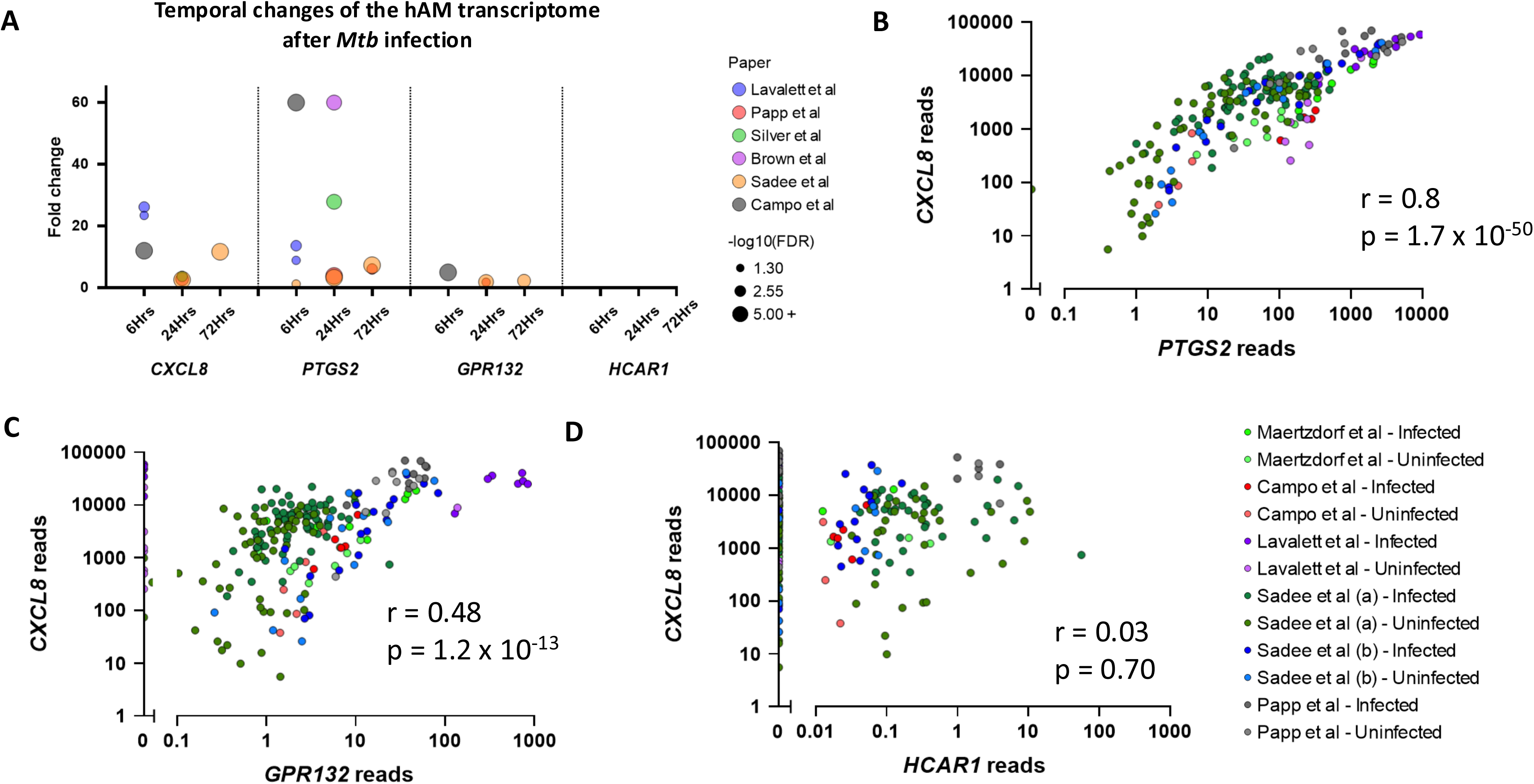
*CXCL8* expression correlates with *PTGS2* and *GPR132* in hAMs infected with *Mycobacterium tuberculosis*. **(A)** An XY scatter plot representing the fold change of specific transcripts after infection with *Mtb* for the specified duration (6h, 24h or 72h), relative to the expression of an uninfected control. The colour represents the source study paper^57–61,92,93^, and the point size represents -log_10_(FDR). *Mtb* H37Rv was used for all studies, except for Lavalett *et al.* where UT127 and UT205 were used instead^57^. Gene expression correlation plots between *CXCL8* and **(B)** *PTGS2*, **(C)** *GPR132*, and **(D)** *GPR81*/*HCAR1*, with the colour representing the source paper and the shade representing the infection (dark) and control (light) groups.

### GPR132 signalling is required for lactate-induced IL-8 secretion in human alveolar macrophages

Next, *in vitro* experiments were designed to confirm the potential role of COX enzymes and/or GPR132 in the secretion of lactate-driven IL-8 in hAMs. First, hAMs were treated with lactate in the presence of the COX-1/2 inhibitor indomethacin, or the specific COX-2 inhibitor NS398. Neither approach reduced the secretion of IL-8 upon lactate treatment, suggesting that COX products do not play a role in this process (Figure 4A). Moreover, to further rule out a role for PGE2 in driving IL-8 secretion in this context, hAMs were treated with increasing concentrations of PGE2, which resulted in decreased, rather than increased, IL-8 production (Figure 4B). Having dismissed PGE2 as a driver of IL-8 secretion in lactate-treated hAMs, the potential role of the specific lactate receptor GPR132 was then explored. We first confirmed that hAMs express GPR132 (Figure 4C). Moreover, lactate signals in a dose-dependent manner through GPR132, and millimolar concentrations are sufficient to induce GPR132-dependent signalling (Figure 4D). Supplementary Figure 3A and 3B show that 9-HODE and SB31 are GPR132 agonists, and that GSK5A (GSK1820795A) is a GPR132 antagonist, in line with previous literature^68^. Antagonising GPR132 with GSK5A prevented lactate-driven IL-8 secretion in hAMs (Figure 4E), and this effect was dose-dependent (Figure 4F). The requirement of GPR132 signalling for IL-8-driven secretion in hAMs was confirmed by using SB-55 (SB-583355) another, structurally distinct^68,69^, GPR132 antagonist (Figure 4E). Thus, GPR132 but not PGE2 is required for lactate-driven IL-8 in hAMs.

**Figure 4.**
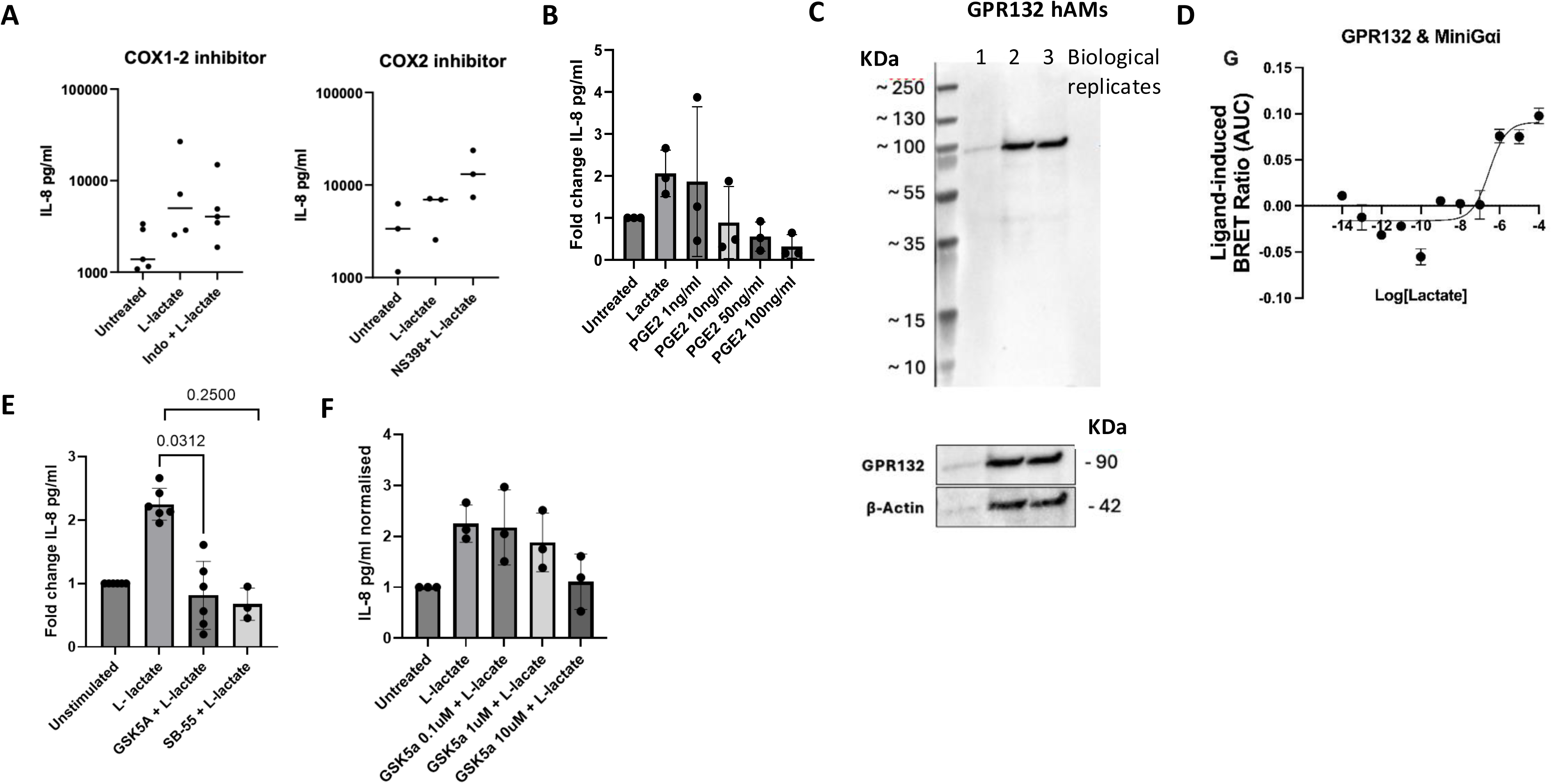
GPR132 signalling is required for lactate-induced IL-8 secretion in human alveolar macrophages. Concentration of IL-8 in hAMs’ supernatants treated with **(A)** Na-lactate (30mM, 48h) and the COX-1 and COX-2 inhibitor indomethacin, or **(B)** increasing concentrations of PGE2. **(C)** Western blot of GPR132 in lysates from hAMs (n=3). **(D)** Lactate concentration-response assay for Mini Gαi recruitment to GPR132-YFP in HEK293 cells (n=3). Concentration of IL-8 in hAMs’ supernatants treated with **(E)** Na-lactate (30mM, 48h) and the GPR132 antagonists GSK5a and SB-55 (both at 10μM) or **(F)** Na-lactate (30mM, 48h) and different concentrations of the GPR132 antagonist GSK5a, measured by ELISA (n=3-6).b

## DISCUSSION

Here, we describe how lactate, the end-product of glycolysis, acts as a metabolic and immune-modulator of hAMs. Specifically, we unravelled the role of lactate in IL-8 production by hAMs. Lactate-driven IL-8 secretion requires the specific lactate transporter MCT1, as well as lipid metabolism reprogramming and GPR132 signalling. Thus, the data presented herein unveil previously unrecognized mechanisms. This dual requirement (lactate transporter and receptor) provides new insights into the role of lactate in hAM function and opens new avenues for therapeutic intervention (Figure 5).

**Figure 5.**
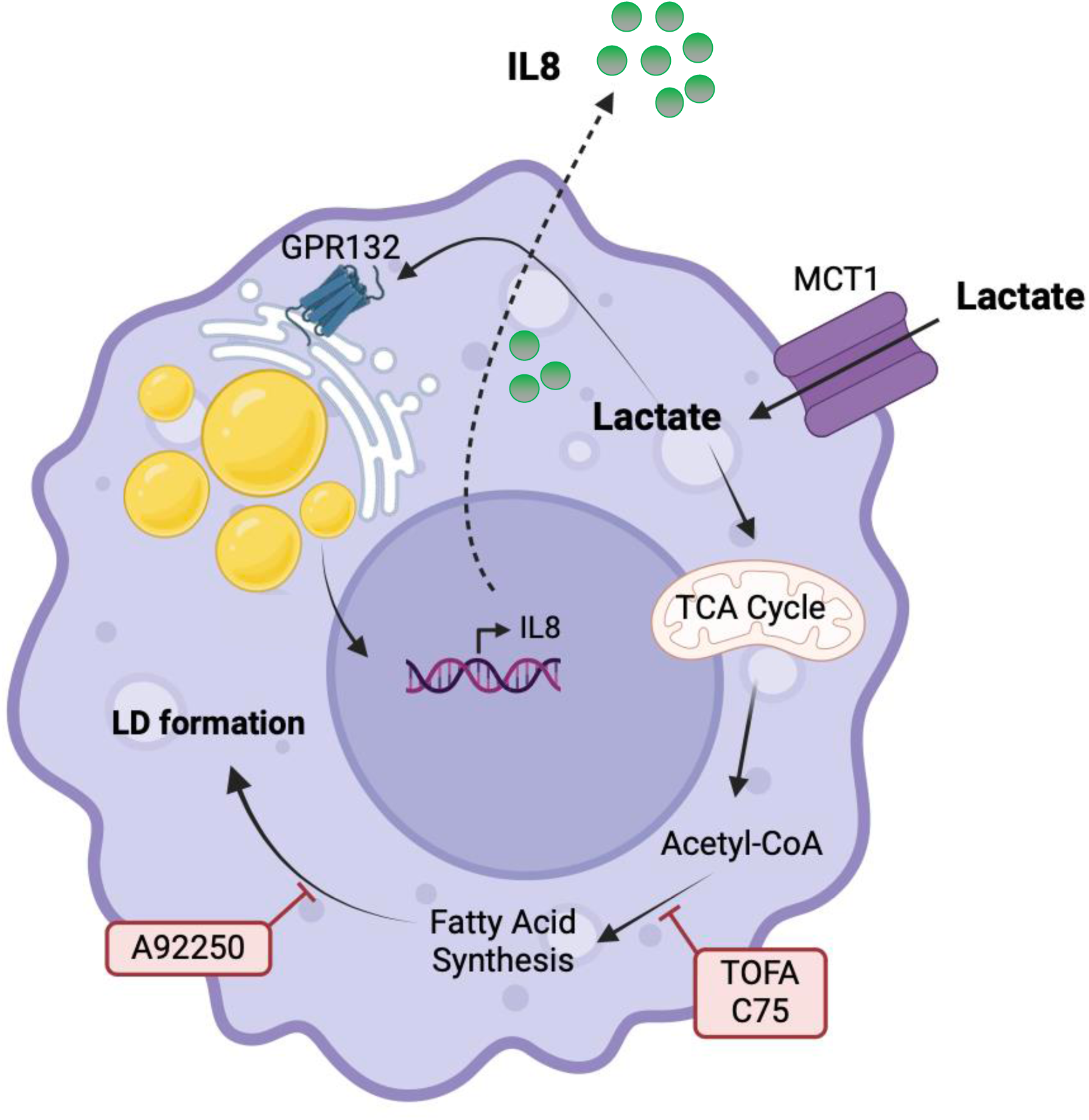
Graphical summary. hAMs uptake environmental lactate via the MCT1 transporter. Lactate signals through GPR132 and promotes lipid synthesis and accumulation, facilitating a foamy macrophage phenotype and IL-8 secretion.

In this study, we investigated lactate-driven IL-8 secretion and the role of two specific lactate transporters (MCT1 and MCT4) in hAMs. Future work should explore the potential contribution of other transporters in the lactate flux in hAMs (e.g. SLC5A12), as well as the mechanisms involved in lactate modulating the secretion of different cytokines and immune-mediators (e.g. IL-6, MIP-1α). Despite abundant evidence of increased glycolysis during lung inflammation^70^, the precise *in vivo* concentration of lactate in the diseased lung remains unknown. Lactate is present at concentrations of 20-50 mM in the lungs from cystic fibrosis patients^29–33^ and tumour fluids^34,35^. Although we show a dose-dependent response to lactate both for IL-8 secretion (Figure 1D) and GPR132 signalling (Figure 4D), it would be useful to determine what is the physiological lactate concentration range in the human lung, both in health and disease.

Lactate fuels the TCA cycle, and its carbons can be used to synthesise acetyl-CoA^25–27^, the precursor for fatty acid synthesis^28^ (Figure 2C and Figure 5), as demonstrated in human CD4+ T cells during chronic inflammation^27^. The association between the foamy macrophage phenotype and IL-8 secretion has been described in atherosclerosis^71,72^ and in a THP-1 model of *Mtb* infection^73^. Interestingly, GPR132 expression has been reported in macrophages from atherosclerosis plaques in mice, rabbits, and humans, with higher levels in macrophages from lipid-rich plaques compared to those from fibrous ones^74^.

Investigation of publicly available gene expression datasets of hAMs infected with *Mtb* revealed a strong correlation between *CXCL8* and both *PTGS2* and *GPR132* (Figure 3B, 3C and 3D). While *in vitro* experiments confirmed the role of GPR132 on lactate-driven IL-8 secretion (Figure 4E and 4F), the obtained results did not support a role for COX enzymes (Figure 4A and 4B). Since COX enzymes - and PGE2 specifically - have been described as drivers of IL-8 production in different cell types and contexts^50–53^, it would be interesting to further study and understand the peculiarities of hAMs, and why PGE2 does not promote IL-8 secretion in their specific context. Interestingly, a study from Standiford and colleagues described PGE2-mediated inhibition of both IL-8 mRNA and protein in LPS-treated hMDMs, and no effects on hAMs^75^. Thus, the interpretation of molecular and cellular processes must account for the specific cellular context, as signalling pathways and functional outcomes are highly dependent on the microenvironment and cell type in which they occur. It would also be interesting to investigate recently identified GPCRs involved in lactate signalling in hAMs (i.e. GPR55^76^, GPR65^77^, GPR109^78^). Lactate signalling through GPR132 involved recruitment of the Gαi subunit (Figure 4D). Further work should help elucidate downstream intracellular mediators (e.g. cAMP, Ca^2+^), as well as the precise location of GPCR signalling (e.g. plasma membrane versus intracellular compartments)^55,79^.

This work focuses on the impact of lactate on a single cell type, namely the hAM. Although beyond the scope of this study, it is important to investigate how lactate shapes the metabolism and function of other lung cell types, including immune (e.g. neutrophils, interstitial macrophages) and non-immune cells (e.g. alveolar epithelial cells, fibroblasts). Furthermore, lactate not only impacts immune cell behaviours but also affects pathogens, serving both as a carbon source and as a modulator of virulence through protein lactylation^64,80,81^. A comprehensive understanding of the impact of lactate on distinct cell types and microorganisms is essential if we are to manipulate lactate sensing and signalling pathways for therapeutic purposes.

Different animal models including mice, rabbits, guinea pigs, cattle and non-human primates (NHP) have been employed to study the immunology of TB^82^. While studies using these models have contributed to a better understanding of TB pathogenesis, none of them accurately recapitulate the features of human TB. *Mtb* has evolved within humans for about 15,000 years and it is a remarkably effective human pathogen^83^. Therefore, using human lung primary cells to model *in vivo* immune responses is paramount and could aid the development of effective new treatments for a range of lung pathologies. Many studies have modelled *Mtb* infection and subsequent immune responses in hMDMs. hMDMs are a useful but limited model, since they do not fully recapitulate the functional and molecular features of primary tissue macrophages. Specifically, differences at the transcriptomic and proteomic levels have been described between hAMs and hMDMs^84–87^. Further differences between these two macrophage subsets have been described at the metabolic level^88^, and in the context of *Mtb* infection^61^. Thus, a real strength of this work is the use of primary hAMs in most experimental settings.

The detailed mechanistic insights into the lactate–IL-8 axis presented here provide a basis for considering new therapeutic strategies, since IL-8 is a key cytokine in several lung diseases, exerting both protective and detrimental effects^6^. Interestingly, modulators of lactate uptake (i.e. MCT1 inhibitors) are being trialled as anti-cancer therapies^89,90^. Considering the global burden of lung diseases and the limited successful therapeutic options for some of them, we urgently need new approaches. Immune therapies have revolutionised the cancer field. Host-directed therapies offer great potential to fight old diseases in a new way. To design them, a comprehensive understanding of specific host immune cellular pathways is essential. The fact that GPR132 is crucial to the lactate-elicited responses here described is noteworthy, since approximately one third of all drugs available today target GPCRs ^91^. This study brings new insights on how lactate, a common molecule in the lung in the context of inflammation and infection, impacts the function of the primary immune cell type in the pulmonary niche (i.e. the hAM). Exploiting this knowledge may aid the design of new therapies for lung diseases.

## MATERIALS AND METHODS

### Lung tissue & blood ethics

The use of lung tissue for laboratory research was approved by the Regional Ethical Approval Committee (REC: 17/WM/0272). Lung tissue was obtained from both male and female patients, non-smokers or long-term ex-smokers (> 5 years) undergoing therapeutic lung cancer surgery at The Queen Elizabeth Hospital (Birmingham, UK). Lung conditions (e.g. COPD, ARDS) were exclusion criteria. All participants gave written informed consent. Healthy, tumour-adjacent tissue was stored in 0.9% saline at 4°C until used for human alveolar macrophage (hAM) isolation.

### PBMC isolation and hMDM differentiation

Human peripheral blood mononuclear cells (PBMCs) were freshly isolated from venous whole blood of healthy donors (15/NW/0079). Whole blood was diluted in PBS (1:1 ratio). 35ml of diluted blood were layered on 12ml of Lymphoprep (Stemcell) in a 50ml falcon tube, which was spun at 800g (0 brake, 0 acceleration) at room temperature for 30 minutes. PBMCs were then isolated from the gradient with a Pasteur pipette into a fresh falcon tube, which was topped up with PBS and spun at 500g, room temperature for 5 minutes. The supernatant was discarded, and the wash step was repeated. PBMCs were then resuspended in warmed in R10 complete media (RPMI 1640 (Sigma) + 10% FCS (Sigma-Aldrich) + 1% Penicillin-Streptomycin (Gibco). Cells were then counted on a haemocytometer and R10 media was added to reach a concentration of 1×10^6^ cells/ml. PBMCs were plated (3×10^6^ cells/3ml in a 6-well plate or 5×10^5^ cells/500uL in a 24-well plate) on Day 0 and incubated at 37oC in 5% CO_2_. On Day 1, the media was changed removing cells in suspension, while the monocyte population remained in the wells due to their adherence capacity. Macrophage colony-stimulating factor (M-CSF; Miltenyi Biotec) was added at 10ng/ml to promote macrophage differentiation. Following a media change on Day 3, PBMCs differentiated into macrophages over 7 days.

### hAM isolation

To isolate alveolar macrophages, 0.9% saline was injected into the airways of the lung tissue using an intravenous giving set and a 25-gauge needle. This caused tissue inflation and facilitated subsequent flushing out of resident lung cells, including alveolar macrophages. Lavage fluid was poured into 50 ml Falcon tubes and spun at 4°C 500g (acceleration 9, brake 9) for 10 minutes. The supernatant was discarded, and the pellets combined and topped up with PBS (Gibco) up to 18 ml. 12 ml of Lymphoprep (Stemcell) were added to an empty Falcon tube to which the pellet-PBS mixture layered. The tube was centrifuged at 800g for 30 minutes (brake 0, acceleration 0), and the mononuclear cells layer extracted. Cells were washed with PBS and centrifuged for 5 minutes at 500g (brake 9, acceleration 9). The supernatant was removed and the cells counted and plated at a density of 500,000 cells/ml in R10 complete media (250μl/well in a 24-well plate). Cells were incubated at 37°C in 5% CO2 overnight. During this time, macrophages will adhere while other cells will remain in suspension; these will be discarded with the media before the addition of fresh media with/without specific treatment/stimulations.

### MCT flow staining

hAMs were detached from the plate using an enzyme-free cell dissociation buffer (Gibco), following manufacturer’s instructions. Cells were washed with PBS (300g for 5 mins) and the supernatant discarded. Cells were fixed (Invitrogen) for 10 minutes at room temperature, and permeabilised (Invitrogen) for 20 minutes and washed with PBS (300g for 5 mins) prior to distribution in BD FACS tubes. These were centrifuged (300g for 5 mins), the supernatants were discarded and the cells were stained), CD68-PE (Miltenyi), MCT1-AF700 (R&D Systems), MCT4-APC (Santa Cruz Biotechnology) or isotype control (IgG2a-AF700, R&D Systems; IgG2a-APC) for 20 minutes at 4°C in the dark. Cells were washed with PBS and resuspended in 300 μL of PBS before using the flow cytometer. Cells were acquired using a LSR Fortessa II (BD Biosciences) flow cytometer, and results were analysed on FlowJo version 10.0.

### hAM treatments

After incubating overnight and changing the media to remove non-adherent cells, hAMs were treated with Na-L-Lactate (Sigma-Aldrich, 30mM unless otherwise stated), Phloretin (Sigma-Aldrich, 25mM), GSK1820795A (GSK5a; GSK, 10μM unless otherwise stated), SB-583355 (SB-55, GSK, 10μM), Indomethacin (Sigma-Aldrich, 10μM), NS398 (Sigma-Aldrich, 100μM), PGE2 (Sigma-Aldrich, 1-100 ng/ml), Oleic Acid (Cayman Chemical Company, 30μM), A922500 (Sigma-Aldrich, 10μM), TOFA (Sigma-Aldrich, 20 μg/ml), C75 (Sigma-Aldrich, 20μM). Lactate was added 1 hour after inhibitors/antagonists. After 48h incubation, supernatants were collected and stored at −80 °C for Luminex (Thermo Fischer Scientific) or IL-8 ELISA (Invitrogen); cells were stained for Oil Red O (see below).

### IL-8 ELISA

Human IL-8 Enzyme-linked Immunosorbent assay (ELISA) kit (Invitrogen) was used following manufacturer’s instructions. Supernatants were thawed at room temperature and centrifuged at 3000g for 5 minutes to remove cell debris. The plate was read using the Synergy HT microplate reader and the Gen5 software.

### Luminex

Human ProcartaPlex Mix&Match 8-plex (IL-1α, IL-1β, IL-1Ra, IL-6, IL-8, MIP-1α, MIP-1β, TNFα; Thermo Fischer Scientific) was used to analyse supernatants from hMDMs and hAMs untreated/treated with lactate, according to manufacturer’s instructions. Measurements were performed using the Bio-Plex 200 system (Bio-Rad).

### ORO staining

A 0.3% stock solution of Oil Red O (ORO; Sigma-Aldrich) in isopropanol was made. hAMs were washed with PBS, then fixed with 4% PFA (Thermo Fischer Scientific) for 30 minutes at room temperature. ORO stock solution was diluted in distilled water at a 2:3 ratio. After fixation, cells were washed with distilled water and incubated with 60% isopropanol for 5 minutes at room temperature. Isopronanol was removed and diluted ORO solution added for 15 minutes at room temperature, after which the ORO solution was removed and cells washed with distilled water until clear. PBS was then added before imaging.

### Microscopy and lipid staining quantification

ToupView software was used to randomly take images of hAMs stained with ORO using a Zeiss AxioVert A1 (Zeiss, Germany) light inverted optical microscope at 10x magnification. Five pictures were randomly taken for each condition. Images were analysed using Image J. For each image, the thirty brightest cells were selected with an annotation tool, and the mean staining of each cell was acquired. The mean staining intensity of the thirty cells was calculated per image, and then the average of the five images was calculated to make a single data point. The data was then put together in GraphPad Prism 10.

### Western blot

hAMs were harvested by detaching cells with an enzyme-free cell dissociation buffer (Gibco) according to the manufacturer’s instructions. Cells were washed twice with ice-cold phosphate-buffered saline (PBS, fisher scientific); the final wash was performed for 10 minutes at 10,000 × g at 4°C. After removing the supernatant, cells were lysed in 200 μL of RIPA buffer (50 mM Tris-HCl, pH 7.4, 150 mM NaCl, 1% NP-40, 0.5% sodium deoxycholate, 0.1% SDS; Sigma-Aldrich) supplemented with 10% protease inhibitor cocktail (Sigma-Aldrich) and incubated on ice for 30 minutes with gentle flicking every 5–10 minutes. Lysates were clarified by centrifugation at 10,000 × g for 10 minutes at 4°C. Supernatants containing soluble proteins were collected and stored at −80°C until analysis.

Protein concentration was determined using the Bradford protein assay kit (Bio-Rad). For SDS-PAGE, 30 μg of total protein per sample was mixed with XT sample buffer (Bio-Rad) and XT reducing agent (Bio-Rad) and heated under protein-specific conditions: GPR132 lysates were incubated at 37 °C for 60 minutes, whereas ACC and pACC lysates were heated at 90 °C for 5 minutes. Samples were briefly centrifuged at 10,000 × g for 30 seconds before loading onto a 4–12% Criterion XT Bis-Tris Protein Gel (Bio-Rad). Electrophoresis was performed at 100 V for 2 hours using distilled water-diluted XT MES running buffer (Bio-Rad). Proteins were transferred to 0.2 μm nitrocellulose membranes using the Trans-Blot Turbo Midi Transfer Pack (Bio-Rad) in the Trans-Blot Turbo Transfer System following manufacturer’s instructions.

Membranes were washed three times with Tris-buffered saline containing 0.1% Tween-20 (TBST, ThermoFisher) and blocked with 5% milk in TBST for 1 hour at room temperature on a shaker (60 rpm). Membranes were incubated overnight at 4°C with primary antibodies: anti-ACC (Cell Signalling Technology, 1:1000), anti-pACC (Cell Signalling Technology, 1:1000), and anti-GPR132 (Abcam, 1:500). After three washes with TBST, membranes were incubated for 1 hour at room temperature with HRP-conjugated secondary antibodies (anti-rabbit IgG-HRP, Cell Signalling Technology, 1:2000). Membranes were washed three additional times with TBST.

Protein bands were visualized using Clarity Western ECL Peroxide and Luminol/Enhancer reagents (Bio-Rad) for 5 minutes at room temperature. Chemiluminescent signals were captured with the ChemiDoc MP Imaging System (Bio-Rad) using exposure times optimized to prevent signal saturation. Band intensities were quantified using ImageJ software and normalised to loading controls (β-actin).

### Analysis of publicly available datasets

For Figure 3, The gene expression and fold change (FC) information was obtained from the GEO-deposited dataset for each paper. The FC was provided directly or calculated from the normalised transcript expression levels. The transcripts per million (TPM)-normalised transcript read values were generated automatically by the NCBI for each GEO-deposited dataset. GraphPad Prism 10.5.0 was used to perform normality tests on the infected and uninfected reads values for each transcript. As these tests failed, the Spearman correlation test was applied. GraphPad Prism 10.5.0. was used for statistical analysis and plotting. For Supplementary Figure 2, Data was obtained from the full Human Lung Cell Atlas (CZ CELLxGENE: Discover)^67^, subset by transcript in human alveolar macrophages to correlate *CXCL8* with *PTGS2*, *GPR132*, and *GPR81*/*HCAR1*, then subset by disease conditions including Normal, COVID-19, Pulmonary Fibrosis, Chronic Obstructive Pulmonary Disease (COPD) and Pneumonia.

### Bioluminescence resonance energy transfer (BRET)

For Figure 4D, Bioluminescence resonance energy transfer (BRET) experiments were performed in Human embryonic kidney 293T (HEK293T) cells as previously described^55^. For miniGsi experiments shown in Supplementary Figure 3, GSK1820795A, SB-583831, 9-HODE, and SKF-9566789 were gifts from GSK (Stevenage, UK). HEK293T cells were seeded at 7.5 × 10⁵ cells/10 cm dish (Greiner Bio-One, Germany) in DMEM with 10% FBS. After 24 hr, cells were transfected with 1 μg GPR132-Nluc (obtained from Genewiz Azenta; Burlington, MA, USA) and 4 μg Venus-tagged miniGsi (gift from Nevin Lambert, Augusta University) using polyethyleneimine (PEI; 1:6 DNA:PEI ratio). White 96-well plates (Greiner Bio-One, Germany) were coated with 50 μL/well poly-D-lysine (20 μg/mL in PBS) for 20 min and washed with PBS. 24 hours post-transfection, cells were resuspended in DMEM supplemented with 10% FBS and plated onto poly-D-lysine-coated white 96-well plates at 1 × 10^4^ cells/well and incubated at 37 °C in 5% CO₂. 48 hr post-transfection, media was replaced with HBSS (pH 7.45; 145 mM NaCl, 5 mM KCl, 10 mM d-glucose, 2 mM sodium pyruvate, 1 mM MgSO₄·7H₂O, 1.7 mM CaCl₂, 1.5 mM NaHCO₃, 10 mM HEPES) and cells treated with GPR132 agonists and antagonists in the presence of 5 μM Furimazine (Promega). After 30 min, BRET measurements were acquired using a PHERAstar FSX microplate reader (BMG Labtech) with a BRET1 filter set (475 ± 30 nm and 535 ± 30 nm) over 30 cycles. BRET ratios were calculated as fluorescence/luminescence and normalised to the SB-583831 response, with 0-100% defined as the basal and maximal ligand response respectively.

### Statistical analysis

Non-parametric paired T-test were employed when comparing two groups. When more than two groups were compared, one-way ANOVA was used. For correlations, as data were not normally distributed a Spearman test was applied. Statistical analyses were performed using GraphPad Prism version 10.

## ACKNOWLEDGEMENTS

We are grateful to all study participants, the wider research team who supported this study from University Hospitals Birmingham NHS Foundation Trust and the University of Birmingham, and to the Flow Cytometry platform at the University of Birmingham. Thank you to Prof Galina Mukamolova (University of Leicester) and Prof Sarah Dimeloe (University of Birmingham) for providing feedback on this manuscript. Thank you to Prof Albert Pol, Dr Marta Bosch and Dr Alba Fajardo for guidance on lipid-related experiments.

## AUTHOR CONTRIBUTIONS

AL conceived the study and obtained funding. AL and CM provided overall guidance. AL, MA and MC designed and performed experiments, analysed and interpreted data. JJ analysed publicly available datasets. LD, JH, HH, COM, AG, SC, LA and SLO performed additional experiments. BN provided human lung clinical samples. DC, RL, DT and AS provided guidance on specific aspects of the project and experimental design. AL and CM prepared the manuscript. All authors contributed to manuscript revision, read and approved the submitted version.

## REFERENCES

1. WHO Tuberculosis Report 2025. World Health Organization. 2025.

2. Agudelo, C. W., Samaha, G. & Garcia-Arcos, I. Alveolar lipids in pulmonary disease. A review. Lipids in Health and Disease vol. 19 (2020).

3. Zazara, D. E., Belios, I., Lücke, J., Zhang, T. & Giannou, A. D. Tissue-resident immunity in the lung: a first-line defense at the environmental interface. Seminars in Immunopathology vol. 44 827–854 (2022).

4. Yoshimura, T., Matsushima, K., Oppenheim, J.J., Leonard, E.J. Neutrophil chemotactic factor produced by lipopolysaccharide (LPS)-stimulated human blood mononuclear leukocytes: partial characterization and separation from interleukin 1 (IL-1). The Journal of Immunology 139(3):788–93 (1987).

5. Cambier, S., Gouwy, M. & Proost, P. The chemokines CXCL8 and CXCL12: molecular and functional properties, role in disease and efforts towards pharmacological intervention. Cellular and Molecular Immunology vol. 20 217–251 (2023).

6. Zhang, F. et al. Neutrophil diversity and function in health and disease. Signal Transduction and Targeted Therapy vol. 9 (2024).

7. Ogger, P. P. & Byrne, A. J. Macrophage metabolic reprogramming during chronic lung disease. Mucosal Immunology vol. 14 282–295 (2021).

8. Neill, L. A. J. O., Kishton, R. J. & Rathmell, J. A guide to immunometabolism for immunologists. Nat Rev Immunol 16, 553–65 (2016).

9. Liang, Q., Wang, Y. & Li, Z. Lipid metabolism reprogramming in chronic obstructive pulmonary disease. Molecular Medicine vol. 31 (2025).

10. Little, I., Bersie, S., Redente, E. F., Mccubbrey, A. L. & Tarling, E. J. Alveolar macrophages: guardians of the alveolar lipid galaxy. Current Opinion in Lipidology vol. 36 153–162 (2025).

11. Kotlyarov, S. Linking Lipid Metabolism and Immune Function: New Insights into Chronic Respiratory Diseases. Pathophysiology vol. 32 (2025).

12. Russell, D. G., Cardona, P. J., Kim, M. J., Allain, S. & Altare, F. Foamy macrophages and the progression of the human tuberculosis granuloma. Nature Immunology vol. 10 943–948 (2009).

13. Agarwal, P., Gordon, S. & Martinez, F. O. Foam Cell Macrophages in Tuberculosis. Frontiers in Immunology vol. 12 (2021).

14. Shim, D., Kim, H. & Shin, S. J. Mycobacterium tuberculosis Infection-Driven Foamy Macrophages and Their Implications in Tuberculosis Control as Targets for Host-Directed Therapy. Frontiers in Immunology vol. 11 (2020).

15. Laval, T., Chaumont, L. & Demangel, C. Not too fat to fight: The emerging role of macrophage fatty acid metabolism in immunity to Mycobacterium tuberculosis. Immunological Reviews vol. 301 84–97 (2021).

16. Jia, Q., Yuan, Q., Chen, X. & Hu, Z. Lactate and Lactylation in Respiratory Diseases: from Molecular Mechanisms to Targeted Strategies. Lung vol. 203 (2025).

17. Soto-Heredero, G., Gómez de las Heras, M. M., Gabandé-Rodríguez, E., Oller, J. & Mittelbrunn, M. Glycolysis – a key player in the inflammatory response. FEBS Journal vol. 287 3350–3369 (2020).

18. Hu, T. et al. Metabolic regulation of the immune system in health and diseases: mechanisms and interventions. Signal Transduction and Targeted Therapy vol. (2024).

19. Llibre, A., Kucuk, S., Gope, A., Certo, M. & Mauro, C. Lactate: A key regulator of the immune response. Immunity vol. 58 535–554 (2025).

20. Liu, Z. & Wang, M. Tranporters of monocarbocylates: characterization and functional roles. Medical research archives. (2015). ISSN 2375-1924. Available at: <https://esmed.org/MRA/mra/article/view/68>. Date accessed: 27 jan. 2026.

21. Cai, T. Q. et al. Role of GPR81 in lactate-mediated reduction of adipose lipolysis. Biochem Biophys Res Commun 377, 987–991 (2008).

22. Liu, C. et al. Lactate inhibits lipolysis in fat cells through activation of an orphan G-protein-coupled receptor, GPR81. Journal of Biological Chemistry 284, 2811–2822 (2009).

23. Chen, P. et al. Gpr132 sensing of lactate mediates tumor-macrophage interplay to promote breast cancer metastasis. Proc Natl Acad Sci U S A 114, 580–585 (2017).

24. Murugan Poongkavithai Vadevoo, S., et al. The macrophage odorant receptor Olfr78 mediates the lactate-induced M2 phenotype of tumor-associated macrophages. Proc Natl Acad Sci USA. 118(37):e2102434118 (2021).

25. Brooks, G. A. Lactate as a fulcrum of metabolism. Redox Biology vol. 35 (2020).

26. Li, X. et al. Lactate metabolism in human health and disease. Signal Transduction and Targeted Therapy vol. 7 (2022).

27. Pucino, V. et al. Lactate Buildup at the Site of Chronic Inflammation Promotes Disease by Inducing CD4+ T Cell Metabolic Rewiring. Cell Metab 30, 1055–1074.e8 (2019).

28. Kayser, O., Averesch, N.J.H. (2025). Fatty Acid Biosynthesis and ABE Metabolism. In: Technical Biochemistry. Springer, Wiesbaden. 10.1007/978-3-658-47121-7_10.

29. C Meyer, K., Amessoudji, A., Hollatz, T. & HS Tsao, F. Lactic acid concentrations in bronchoalveolar lavage fluid correlate with neutrophil influx in cystic fibrosis. Archives of Biomedical and Clinical Research 1, (2019).

30. Wolak, J. E., Esther, C. R. & O’Connell, T. M. Metabolomic analysis of bronchoalveolar lavage fluid from cystic fibrosis patients. Biomarkers 14, 55–60 (2009).

31. Haeger, S. et al. The bronchoalveolar lavage dilution conundrum: an updated view on a long-standing problem. Am J Physiol Lung Cell Mol Physiol. 327(5):L807–L813 (2024).

32. Baldesi, O. et al. Bacterial ventilator-associated pneumonia: Bronchoalveolar lavage results are not influenced by dilution. Intensive Care Med 35, 1210–1215 (2009).

33. Drusano, G. L. et al. Dilution factor of quantitative bacterial cultures obtained by bronchoalveolar lavage in patients with ventilator-associated bacterial pneumonia. Antimicrob Agents Chemother 62, (2018).

34. Colegio, O. R. et al. Functional polarization of tumour-associated macrophages by tumour-derived lactic acid. Nature 513, 559–563 (2014).

35. Feng, Q. et al. Lactate increases stemness of CD8 + T cells to augment anti-tumor immunity. Nat Commun 13, (2022).

36. Ingersoll, M. A., Platt, A. M., Potteaux, S. & Randolph, G. J. Monocyte trafficking in acute and chronic inflammation. Trends in Immunology vol. 32 470–477 (2011).

37. Bickel, M. The role of interleukin-8 in inflammation and mechanisms of regulation. J Periodontol. 64(5)):456–60 (1993).

38. Cesta, M. C. et al. The Role of Interleukin-8 in Lung Inflammation and Injury: Implications for the Management of COVID-19 and Hyperinflammatory Acute Respiratory Distress Syndrome. Frontiers in Pharmacology vol. 12 (2022).

39. Patel, M. S., Jomain-Baum, M., Ballard, F. J. & Hanson, R. W. Pathway of carbon flow during fatty acid synthesis from lactate and pyruvate in rat adipose tissue. J Lipid Res 12, 179–191 (1971).

40. Rath, E. A., Beloff-Chain, A. & Hems, D. A. Contribution of Lactate Carbon to Fatty Acid Synthesis in Adipose Tissue of Normal and Genetically Obese (Oblob) Mice. (1975).

41. Robertson, J. P., Faulkner, A. & Vernon, R. G. L-Lactate as a source of carbon for fatty acid synthesis in adult and foetal sheep. Biochirnica et Biophysics Acta vol. 665 (3).511–8 (1981)

42. Qi, J. et al. The use of stable isotope-labeled glycerol and oleic acid to differentiate the hepatic functions of DGAT1 and −2. J Lipid Res 53, 1106–1116 (2012).

43. Castillo-Quan, J. I. et al. An Antisteatosis Response Regulated by Oleic Acid through Lipid Droplet-Mediated ERAD Enhancement. Sci Adv. 9(1):eadc8917 (2023).

44. An, H. et al. New Insights into AMPK, as a Potential Therapeutic Target in Metabolic Dysfunction-Associated Steatotic Liver Disease and Hepatic Fibrosis. Biomolecules and Therapeutics vol. 33 18–38 (2025).

45. Lee, M. et al. Phosphorylation of Acetyl-CoA Carboxylase by AMPK Reduces Renal Fibrosis and Is Essential for the Anti-Fibrotic Effect of Metformin. Journal of the American Society of Nephrology 29(9):p 2326–2336 (2018).

46. Walther, T. C. & Farese, R. V. Lipid droplets and cellular lipid metabolism. Annu Rev Biochem 81, 687–714 (2012).

47. Melo, R. C. N. et al. Lipid bodies in inflammatory cells: Structure, function, and current imaging techniques. Journal of Histochemistry and Cytochemistry vol. 59 540–556 (2011).

48. Martínez-Colón, G. J. & Moore, B. B. Prostaglandin E2 as a Regulator of Immunity to Pathogens. Pharmacology and Therapeutics vol. 185 135–146 (2018).

49. An, Y., Yao, J. & Niu, X. The Signaling Pathway of PGE2and Its Regulatory Role in T Cell Differentiation. Mediators of Inflammation vol. 2021 (2021).

50. Denizot, Y., Godard, A., Raher, S., Trimoreau, F. & Praloran, V. Lipid mediators modulate the synthesis of inertelukin 8 by human bone marrow stromal cells. Cytokine. 11(8)606–10 (1999).

51. Yu, Y. & Chadee, K. Prostaglandin E3 stimulates IL-8 gene expression in human colonic epithelial cells by a posttranscriptional mechanism. The Journal of Immunology 161(7)3746–52. (1998).

52. Caristi, S. et al. Prostaglandin E2 induces interleukin-8 gene transcription by activating C/EBP homologous protein in human T lymphocytes. Journal of Biological Chemistry 280, 14433–14442 (2005).

53. Srivastava, V., Dey, I., Leung, P. & Chadee, K. Prostaglandin E 2 modulates IL-8 expression through formation of a multiprotein enhanceosome in human colonic epithelial cells. Eur J Immunol 42, 912–923 (2012).

54. Certo, M., Llibre, A., Lee, W. & Mauro, C. Understanding lactate sensing and signalling. Trends in Endocrinology and Metabolism vol. 33 722–735 (2022).

55. O’Brien, S. L. et al. Intracrine FFA4 signaling controls lipolysis at lipid droplets. Nat Chem Biol. 22(1):109–119 (2025).

56. Calebiro, D., Miljus, T. & O’Brien, S. Endomembrane GPCR signaling: 15 years on, the quest continues. Trends in Biochemical Sciences vol. 50 46–60 (2025).

57. Lavalett, L., Ortega, H. & Barrera, L. F. Human Alveolar and Splenic Macrophage Populations Display a Distinct Transcriptomic Response to Infection With Mycobacterium tuberculosis. Front Immunol 11, (2020).

58. Papp, A. C. et al. AmpliSeq transcriptome analysis of human alveolar and monocyte-derived macrophages over time in response to Mycobacterium tuberculosis infection. PLoS One 13, (2018).

59. Maertzdorf, J. et al. Mycobacterium tuberculosis Invasion of the Human lung: First contact. Front Immunol 9, (2018).

60. Sadee, W., et al. Human alveolar macrophage response to Mycobacterium tuberculosis: immune characteristics underlying large inter-individual variability. Commun Biol 8, (2025).

61. Campo, M. et al. Human Alveolar and Monocyte-Derived Human Macrophage Responses to Mycobacterium tuberculosis. The Journal of Immunology 213, 161–169 (2024).

62. Shi, L., Eugenin, E. A. & Subbian, S. Immunometabolism in tuberculosis. Frontiers in Immunology vol. 7 (2016).

63. Hackett, E. E. & Sheedy, F. J. An Army Marches on Its Stomach: Metabolic Intermediates as Antimicrobial Mediators in Mycobacterium tuberculosis Infection. Front Cell Infect Microbiol 10, 446 (2020).

64. Llibre, A. et al. Lactate cross-talk in host-pathogen interactions. Biochemical Journal vol. 478:3157–3178 (2021).

65. Zhang, Y. et al. Enhanced interleukin-8 release and gene expression in macrophages after exposure to Mycobacterium tuberculosis and its components. Journal of Clinical Investigation 95, 586–592 (1995).

66. Lyons, M. J., Yoshimura, T. & McMurray, D. N. Interleukin (IL)-8 (CXCL8) induces cytokine expression and superoxide formation by guinea pig neutrophils infected with Mycobacterium tuberculosis. Tuberculosis 84, 283–292 (2004).

67. Abdulla, S. et al. CZ CELLxGENE Discover: A single-cell data platform for scalable exploration, analysis and modeling of aggregated data. Nucleic Acids Res 53, D886–D900 (2025).

68. Foster, J. R. et al. N-Palmitoylglycine and other N-acylamides activate the lipid receptor G2A/GPR132. Pharmacol Res Perspect 7, (2019).

69. Hernandez-Olmos, V. et al. Development of a Potent and Selective G2A (GPR132) Agonist. J Med Chem 67, 10567–10588 (2024).

70. Michaeloudes, C. et al. Role of Metabolic Reprogramming in Pulmonary Innate Immunity and Its Impact on Lung Diseases. Journal of Innate Immunity vol. 12 31–46 (2020).

71. Wang, N. et al. Interleukin 8 Is Induced by Cholesterol Loading of Macrophages and Expressed by Macrophage Foam Cells in Human Atheroma. J Biol Chem 271(15)8837–42(1996).

72. Liu, Y., Hultén, L. M. & Wiklund, O. Macrophages isolated from human atherosclerotic plaques produce IL-8, and oxysterols may have a regulatory function for IL-8 production. Arterioscler Thromb Vasc Biol 17, 317–323 (1997).

73. Sanjurjo, L. et al. The scavenger protein apoptosis inhibitor of macrophages (AIM) potentiates the antimicrobial response against Mycobacterium tuberculosis by enhancing autophagy. PLoS One 8, (2013).

74. Rikitake, Y. et al. Expression of G2A, a receptor for lysophosphatidylcholine, by macrophages in murine, rabbit, and human atherosclerotic plaques. Arterioscler Thromb Vasc Biol 22, 2049–2053 (2002).

75. Standiford, T.J. et al. Regulation of Human Alveolar Macrophage and Blood Monocyte-derived Interleukin-8 by Prostaglandin E_2_ and Dexamethasone. Am J Respir Cell Mol Biol. 6(1):75–81 (1992).

76. Sgrignani, G. et al. GPR55 senses lactate to sustain motility in prostate cancer cells. Mol Cell Biochem 480, 5197–5204 (2025).

77. Yan, C. et al. GPR65 sensing tumor-derived lactate induces HMGB1 release from TAM via the cAMP/PKA/CREB pathway to promote glioma progression. Journal of Experimental and Clinical Cancer Research 43, (2024).

78. Ahmed, K., Tunaru, S. & Offermanns, S. GPR109A, GPR109B and GPR81, a family of hydroxy-carboxylic acid receptors. Trends Pharmacol Sci 30, 557–562 (2009).

79. Calebiro, D. et al. Persistent cAMP-signals triggered by internalized G-protein-coupled receptors. PLoS Biol 7, (2009).

80. Wang, Y. et al. Post-translational toxin modification by lactate controls Staphylococcus aureus virulence. Nature Communications 15, (2024).

81. Zhang, C. et al. Deciphering novel enzymatic and non-enzymatic lysine lactylation in Salmonella. Emerg Microbes Infect 14, (2025).

82. Williams, A. & Orme, I. M. Animal Models of Tuberculosis: An Overview. Microbiol Spectr 4, (2016).

83. Daniel, T. M. The history of tuberculosis. Respir Med 100, 1862–1870 (2006).

84. Li, J. et al. cDNA microarray analysis reveals fundamental differences in the expression profiles of primary human monocytes, monocyte-derived macrophages, and alveolar macrophages. J Leukoc Biol 81, 328–335 (2007).

85. Groot-Kormelink, P. J., Fawcett, L., Wright, P. D., Gosling, M. & Kent, T. C. Quantitative GPCR and ion channel transcriptomics in primary alveolar macrophages and macrophage surrogates. BMC Immunol 13, (2012).

86. Tomechko, S. E. et al. Proteomic and bioinformatics profile of paired human alveolar macrophages and peripheral blood monocytes. Proteomics 15, 3797–3805 (2015).

87. Jin, M., Opalek, J. M., Marsh, C. B. & Wu, H. M. Proteome comparison of alveolar macrophages with monocytes reveals distinct protein characteristics. Am J Respir Cell Mol Biol 31, 322–329 (2004).

88. Pereverzeva, L. et al. Human alveolar macrophages do not rely on glucose metabolism upon activation by lipopolysaccharide. Biochim Biophys Acta Mol Basis Dis 1868, (2022).

89. Benyahia, Z. et al. In vitro and in vivo characterization of mct1 inhibitor azd3965 confirms preclinical safety compatible with breast cancer treatment. Cancers (Basel) 13, 1–25 (2021).

90. Halford, S. et al. A Phase I Dose-Escalation Study of AZD3965, an Oral Monocarboxylate Transporter 1 Inhibitor, in Patients with Advanced Cancer. Clinical Cancer Research 29, 1429–1439 (2023).

91. Lorente, J.S., et al. GPCR drug discovery: new agents, targets and indications. Nat Rev Drug Discov. 24, 458–479 (2025).

92. Brown, K., et al. *Mycobacterium tuberculosis* induces Warburg Metabolism in Human Alveolar Macrophages: A Transcriptomic Analysis. Am J Respir Cell Mol Biol. 72(5):597–602 (2025).

93. Silver, R. F. et al. Human alveolar macrophage gene responses to mycobacterium tuberculosis strains H37Ra and H37Rv. Am J Respir Cell Mol Biol 40, 491–504 (2009).

