## Supplementary figures and images for "Lactate promotes IL-8 secretion in human alveolar macrophages through GPR132 and lipid metabolic reprogramming"

### Supplementary Figures

**Supplementary Figure 1.**

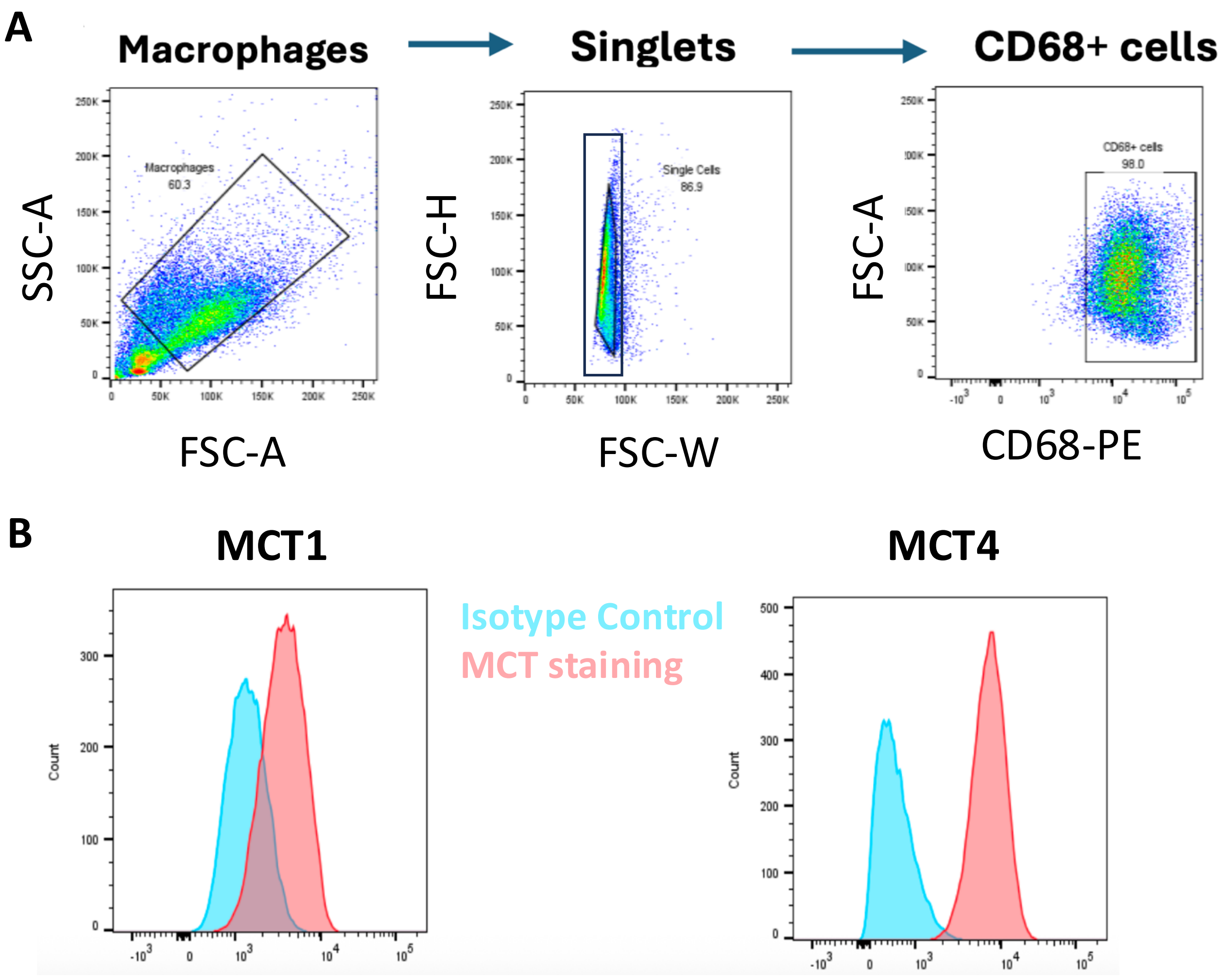

#### Supplementary Figure 2.

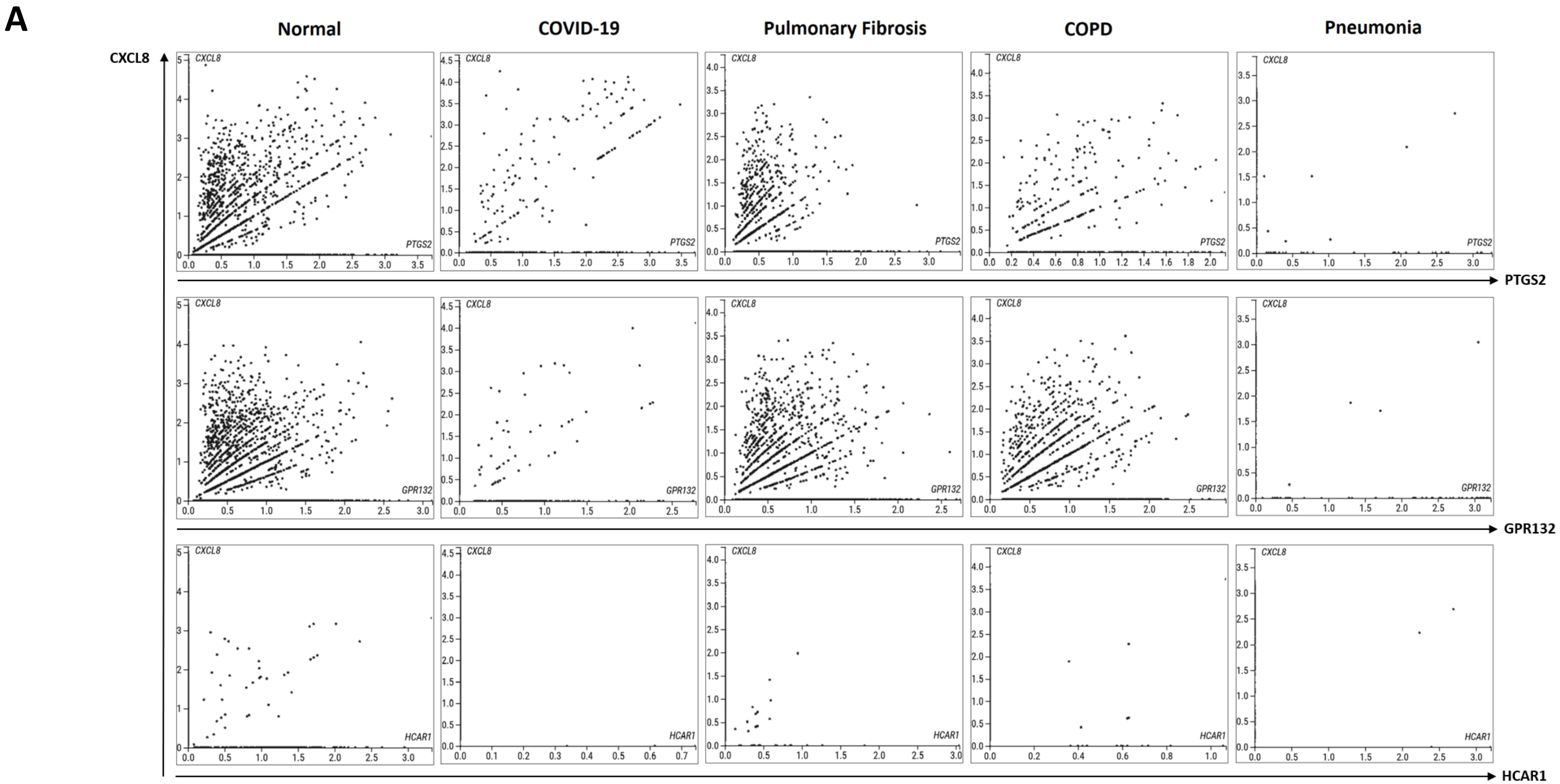

**Supplementary Figure 3.**

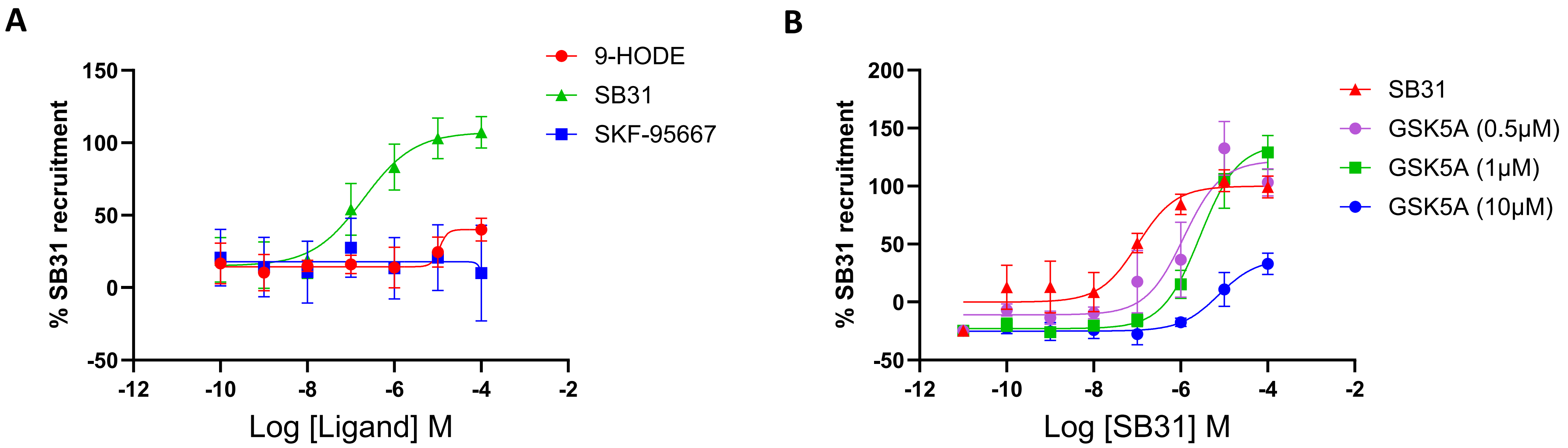
